# Modelling Protein-Glycan Interactions with HADDOCK

**DOI:** 10.1101/2024.07.31.605986

**Authors:** Anna Ranaudo, Marco Giulini, Angela Pelissou Ayuso, Alexandre M. J. J. Bonvin

## Abstract

The term glycan refers to a broad category of molecules composed of monosaccharides units linked to each other in a variety of ways, whose structural diversity is related to different functions in living organisms. Among others, glycans are recognized by proteins with the aim of carrying information and for signalling purposes. Determining the three-dimensional structures of protein-protein-glycan complexes is essential both for the understanding of the mechanisms glycans are involved in, and for applications such as drug design. In this context, molecular docking approaches are of undoubted importance as complementary approaches to experiments. In this study, we show how HADDOCK can be efficiently used for the prediction of protein-glycan complexes. Using a benchmark of 89 complexes, starting from their bound or unbound forms, and assuming some knowledge of the binding site on the protein, our protocol reaches a 70% and 40% top 5 success rate on bound and unbound datasets, respectively. We show that the main limiting factor is related to the complexity of the glycan to be modelled and the associated conformational flexibility.

## Introduction

Glycans are organic compounds with a polymeric structure consisting of monosaccharides, small building blocks covalently linked to each other in various ways through glycosidic bonds, in linear or branched arrangements. Depending on the number of monosaccharides units, glycans are named disaccharides (2 units), oligosaccharides (3-10 units), polysaccharides (> 10 units)^1^. For simplicity, we use here the term “glycan” for referring to compounds of any number of monosaccharide units.

Glycans can form complex structures. This complexity already lies in monosaccharides themselves, having a high degree of intrinsic chemical variability. The second source of complexity arises from the way monosaccharides are linked to each other, as each glycosidic bond can form two possible stereoisomers at the anomeric carbon, i.e., the carbon whose asymmetric center is formed upon the cyclization of the monosaccharide. Regioisomers may also exist because of the many hydroxyl groups, which allow two monosaccharides to be linked in several ways^2^. Monosaccharides, having the ability to create more than two glycosidic bonds, can give rise to branched chains. The frequent occurrence of branched patterns, as opposed to the linear patterns typically found in most peptides and oligonucleotides, contributes to the complexity of glycans’ structural landscape. Besides the very large amount of glycans that can be built starting from a few monosaccharides, glycans show large conformational variability at room temperature, due to the low torsional energy barriers around the glycosidic bonds^3^.

Glycans are ubiquitously found in living organisms, where they can be free-standing or linked to proteins and lipids, giving rise to glycoproteins and glycolipids, respectively. They are involved in a variety of biological processes that can be classified in three main categories^4^. First, they can play structural roles, by either assisting in creating extracellular scaffolds, such as cell walls and matrices, or by being involved in proteins folding and function. Second, they can serve as crucial sources for energy metabolism. Third, they can play the role of information carriers, being recognized by Glycan Binding Proteins (GBPs). Glycans can bind proteins through non-covalent, reversible interactions, for signalling purposes, initiating a range of biological processes in both plants and animals^5^. The role of glycans has been recently highlighted in the context of the well-known SARS-CoV-2 spike protein. This protein, which enters host cells by connecting to the angiotensin-converting enzyme (ACE2), is surrounded by a layer of glycans which hides it from the immune system. Casalino et al.^6^ showed that specific glycans play a crucial role in the movement and structure of the part of the spike protein that binds to ACE2. Their removal results in diminished binding to ACE2.

It is evident that understanding the way glycans interact with proteins is of crucial importance. Experimental methods such as X-ray crystallography and cryo-electron microscopy face challenges when dealing with glycans because of their intrinsic flexibility and heterogeneity. Computational approaches, such as molecular docking, can provide an alternative, less expensive and faster way of generating three-dimensional (3D) models of protein-glycan complexes.

Despite the growing attention that is being given to the carbohydrates field^7,8^, the modelling of protein-glycan complexes by docking has only received limited attention compared to that of other biomolecular complexes. A few protocols have been developed in the past years, such as GlycanDock based on Rosetta^9^, a heparin-specific protocol based on PIPER/ClusPro^10^, VinaCarb^11^ GlycoTorch Vina^12^, RosettaCarbohydrate^13^, and ATTRACT^14^. Recently, a new version of Alphafold^15^, Alphafold3^16^, that can handle glycosylated-proteins has been published. It however does not allow to model protein glycan complexes. State-of-art protein-ligand docking software encounter challenges in addressing the conformational variability of glycans, as they are usually developed for dealing with small, more rigid molecules^9^.

In this study, we use HADDOCK^17,18^, to address the protein-glycan interaction prediction problem. HADDOCK is an information-driven docking approach that can harvest knowledge about binding sites to drive the docking process (see below). Note that HADDOCK^18^ has already been applied to glycans modelling^19,20^, but, to date, without an exhaustive benchmarking. Here we first benchmark the baseline performance of HADDOCK in predicting the 3D structures of protein-glycan complexes on a bound dataset composed of 89 high-resolution experimental complexes from the Protein Data Bank (PDB)^21^ assuming an ideal scenario in which the binding interfaces on both protein and glycan are known, in order to drive the docking process. A protocol is then proposed to deal with a realistic scenario, where the bound conformations of the partners are unknown. The GLYCAM-Web webserver^22,23^ is used for the generation of glycans unbound structures, while the unbound protein structures are retrieved from the PDB. Finally, to address the conformational flexibility challenge of glycans, we assess whether providing an ensemble of glycan conformations generated through a short conformational sampling carried out with HADDOCK prior to the docking process can improve the performance of the protocol, next to the standard semi-flexible refinement of the interface.

## Methods

### Benchmark dataset preparation

HADDOCK’s^17,18^ performance in reproducing the binding geometries of protein-glycan complexes was evaluated by exploiting an adapted version of the dataset provided in GlycanDock^9^. It is composed of 109 experimentally determined high-resolution (< 2.0 Å) protein-glycan complexes collected from the PDB. By discarding the entries containing glycans not yet supported by HADDOCK (refer to https://rascar.science.uu.nl/haddock2.4/library for a list of supported glycans), a dataset of 89 complexes was obtained, which will be referred to as *bound dataset* from now on. The protein receptors in this dataset include 8 antibodies, 21 carbohydrate-binding modules, 18 enzymes, 27 lectins or glycan binding proteins (GBP) and 15 viral glycan binder proteins. The length of the glycans ranges from 2 to 7 monosaccharides units, 72 of which are linear glycans and the remaining 17 branched ones. This structural diversity will be considered in the analysis of the docking performance.

With the aim of evaluating HADDOCK’s performance in a realistic unbound docking scenario, a subset of 55 complexes (out of 89) was defined for which unbound protein forms were available in the PDB. The glycans’ unbound conformations were generated with the GLYCAM-Web webserver^22,23^. This dataset of 55 unbound conformations of both proteins and glycans will be referred to as *unbound dataset* from now on. It contains 47 linear and 8 branched glycans; 25 glycans are composed by three or less monosaccharides units while 30 have more than three units. For the evaluation of HADDOCK’s performance on the *unbound dataset*, all the complexes including glycans made up by three or less monosaccharide units are treated together and referred to with the label **SL-SB** (short linear-short branched). For the larger bound dataset, linear (**SL**) and branched (**SB**) short glycans are analysed separately. The complexes including glycans composed by more than three units (**L** for long) are divided into linear (**LL**, 23 cases) and branched (**LB**, 7 cases). Glycans and protein structures were prepared for docking as described in the <u>Supplementary Text S1</u>. Details on the two datasets are reported in <u>Supplementary Table S1</u>. All HADDOCK-ready bound and unbound conformations, together with restraints, HADDOCK configuration files and analysis scripts, are provided in the following GitHub repository (https://github.com/haddocking/protein-glycans).

### Docking with HADDOCK

Docking calculations were performed with HADDOCK3 (https://github.com/haddocking/haddock3, DOI:10.5281/zenodo.10527751), the new, modular version of the well-established HADDOCK 2.X software^24^. The original HADDOCK protocol consists of three successive steps: i) full randomization of the orientations and docking by rigid-body energy minimization, ii) semi-flexible refinement by simulated annealing in torsion angle space during which the interfaces are considered as flexible, iii) final refinement, either by energy minimization (current default) or by a short molecular dynamics simulation in explicit solvent. HADDOCK3 overcomes this rigid workflow structure as its constituent modules can be freely combined and interchanged by the user.

HADDOCK’s scoring function (HS) includes the intermolecular electrostatic (E_elec_) and van der Waals (E_vdW_) energies (calculated with the OPLS force field^25^), an empirical desolvation energy term (E_desolv_)^26^, the buried surface area (EBSA) and the Ambiguous Interactions Restraints energy (E_air_) (see below). The combination and weights of those terms depend on the stage of the protocol. In the present study, we compare the default scoring function at the rigid body stage (Eq. 1) with one in which the weight of the van der Waals energy term was increased from 0.01 to 1.0 (Eq. 2) as recommended for small molecule docking with HADDOCK^27^. The scoring function in Eq. 2 will be referred to with the name *vdW*, to be distinguished from the *default* one. For the flexible refinement stage, the default scoring function is used (Eq. 3).

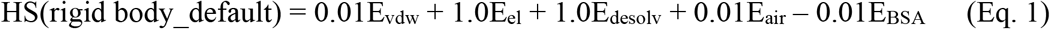

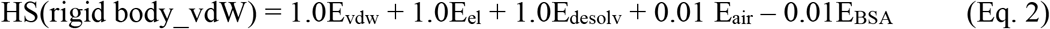

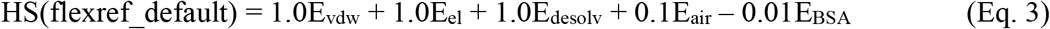

Experimental or predicted information about binding sites can be introduced as restraints for guiding the docking process. Interface information is typically introduced as Ambiguous Interaction Restraints (AIRs)^18^ which are defined from lists of residues divided into two groups: active and passive. Active residues are those of central importance for the interaction. They are restrained to be part of the interface throughout the docking and refinement process; if an active residue is not part of the interface in a model it generates a restraint energy (and corresponding forces). Passive residues are those that could contribute to the interaction; if such a residue is not in the interface of a given model there is no restraint energy generated.

In this work, two scenarios in terms of AIRs were considered: i) true-interface scenario (**ti-aa**), where active residues, corresponding to the interface residues within 3.9 Å^28,29^ from the partner, are defined for both the protein and the glycan; ii) true-interface-protein – full glycan passive scenario (**tip-ap**), where active residues are still defined for the protein interface, but all residues of the glycan are listed as passive. By default, HADDOCK randomly discards 50% of the defined AIRs for each docking model. This is done to account for possible wrong information (false positives) in the experimental data.

### Docking protocols for bound and unbound datasets

Two different protocols were used for the *bound* and *unbound* datasets. For the *bound* dataset a workflow consisting of the following six modules was defined for running HADDOCK3 (details in <u>Supplementary Table S2</u> and corresponding config file available in the GitHub repository):

1. `topoaa`: creation of the topologies of the two partners
2. `rigidbody`: AIR-driven generation of rigid body models
3. `caprieval`: models quality analysis (see below)
4. `rmsdmatrix`: calculation of the RMSD matrix between all the models, based on either all the interface residues (when **ti-aa** AIRs are used) or the protein interface residues and the whole glycan (when **tip-ap** AIRs are used)
5. `clustrmsd`: RMSD-based agglomerative hierarchical clustering of the models using the average linkage criterion and a distance cutoff of 2.5 Å. Only clusters containing four or more models are evaluated.
6. `caprieval`: cluster-based evaluation of the quality of the models

As the starting structures are already in their bound conformations no flexible refinement was performed.

For the *unbound* dataset (details in <u>Table S3</u> and corresponding config file available in the GitHub repository), the workflow consisted of the following twelve modules:

1. `topoaa`: creation of the topologies of the two partners during which any missing atoms are automatically added
2. `rigidbody`: AIR-driven generation of rigid body docking models (1000 by default) with increased sampling (200 per conformation) when starting from an ensemble of conformations
3. `caprieval`: models quality analysis (see below)
4. `rmsdmatrix`: calculation of the RMSD matrix between all the models, based on either all the interface residues (when **ti-aa** AIRs are used) or the protein interface residues and the whole glycan (when **tip-ap** AIRs are used)
5. `clustrmsd`: RMSD-based agglomerative hierarchical clustering of the models using the average linkage criterion. Here 50 (150 when the ensemble of glycans conformations is used) clusters are created.
6. `seletopclusts`: selection of the top 5 models of the existing clusters
7. `caprieval`: cluster-based evaluation of the quality of the models
8. `flexref`: semi-flexible refinement through a simulated annealing protocol in torsion angle space in which first side-chains and then side-chains and backbone of interface residues are treated as flexible
9. `caprieval`: models quality analysis
10. `rmsdmatrix`: calculation of the RMSD matrix between all the models, as in point 4)
11. `clustrmsd`: RMSD-based clustering of the models as in point 4), but here using a distance cutoff of 2.5 Å as in the bound scenario.
12. `caprieval`: cluster-based evaluation of the quality of the models

The workflow for the *unbound* dataset with the RMSD clustering of the rigid body models followed by semi-flexible refinement, is schematically represented in Figure 1.

**Figure 1.**
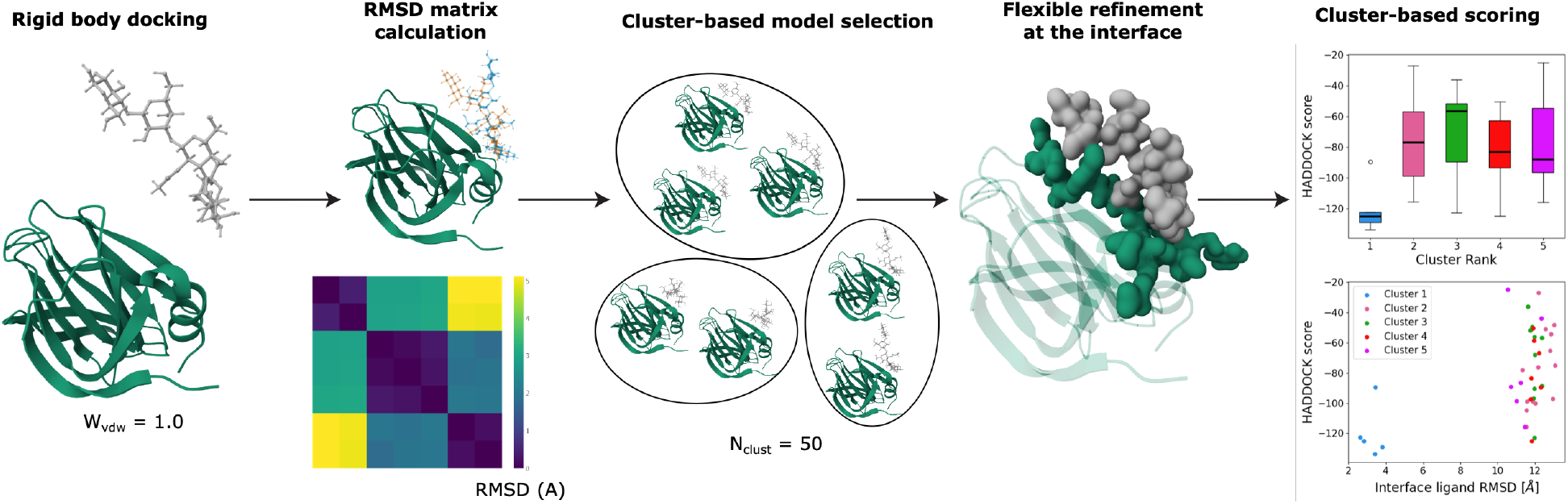
Schematic representation of the docking protocol for the unbound dataset. First, rigid body docking is performed. The models are then clustered based on RMSD. The best scoring models of each cluster are then subjected to a flexible refinement (interface) and the resulting models are again clustered and analysed.

### Model quality assessment

The quality of the models was evaluated using the interface-ligand RMSD (IL-RMSD) with respect to the experimental structures. We didn’t use the Fraction of Native Contacts (Fnat) as its values are typically quite high irrespectively of the pairwise orientation between the molecules and thus unable to highlight small conformational differences, especially for small glycans. The IL-RMSD is calculated by first superimposing the model onto the reference structure using the backbone atoms of the protein interface residues and then calculating the RMSD on the heavy atoms of the oligosaccharide. This is motivated by the fact that the protein interface is larger compared to the glycan interface and would dominate the RMSD calculation if the standard CAPRI^30^ interface RMSD (I-RMSD) metric was used. The IL-RMSD gives a better measure of variations in the position of the ligand compared to the I-RMSD. The cut-offs used to define the quality of the models based on IL-RMSD are:

▪ high-quality models: IL-RMSD ≤ 1.0 Å
▪ medium-quality models: IL-RMSD ≤ 2.0 Å
▪ acceptable-quality models: IL-RMSD ≤ 3.0 Å
▪ near acceptable-quality models: IL-RMSD ≤ 4.0 Å

<u>Supplementary Figure S1</u> shows examples of docking models falling into these four categories for short and long linear glycans and for branched glycans, highlighting how the near acceptable threshold is only appropriate when dealing with long glycans, being too permissive for short monosaccharide chains.

HADDOCK3’s performance is evaluated by calculating success rates (SR) defined as the fraction of complexes having at least one high-, medium-, acceptable-, or near-acceptable-quality model among the top N models ranked according to the HADDOCK score. The cluster-based SR is also calculated by considering the top 4 scoring models within each cluster and assigning a given quality to a cluster if any of the top 4 member of the cluster reaches the corresponding quality cutoff. Note that the clustering step after rigid body docking considers the 5 top models of each cluster.

### Glycans conformational sampling

Conformational sampling of the glycans was carried out with the HADDOCK3 water refinement module (`mdref`^24^), starting from the models generated with the GLYCAM-Web webserver^22,23^. Different scenarios were tested in terms of number of molecular dynamics integration steps and number of models generated (each simulation starting with a different random seed). The RMSD of the generated models with respect to the bound glycan conformation was calculated with the `rmsdmatrix` module. The generated models were clustered using RMSD-based hierarchical clustering^31,32^ requesting 20 clusters. The centres of each cluster were then used to define an ensemble of starting unbound conformations for docking. Further details of the conformational sampling can be found in <u>Supplementary Text S2</u>.

## Results and Discussion

This section is structured as follows. First, the impact of the rigid body scoring function on HADDOCK3’ performance is discussed based on docking calculations performed on the *bound* dataset. The performance of the docking is then analysed considering structural features of the glycans (length and branching) and the definition of the AIRs. We then focus on the more realistic scenario of *unbound docking*, assessing the best way of selecting the rigid body models to be refined and the impact of the flexible refinement on the quality of the final models. Finally, impact and limitations of using an ensemble of glycans as starting point for the docking are discussed.

### Bound docking performance and impact of the rigid body scoring function in the ranking of models

We first assessed the accuracy of the rigid body scoring function in ranking the generated models, comparing the *default* scoring function (Eq. 1) to the one recommended for small ligands, in which the van der Waals energy weight is increased from 0.01 to 1.0 (*vdW* scoring function, Eq. 2). This was assessed by running docking calculations on the *bound dataset* with true interface restraints (**ti-aa** AIRs). A comparison of success rates (SR) obtained with the *default* (Eq. 1) and *vdW* (Eq. 2) scoring functions is shown in Figure 2.

**Figure 2.**
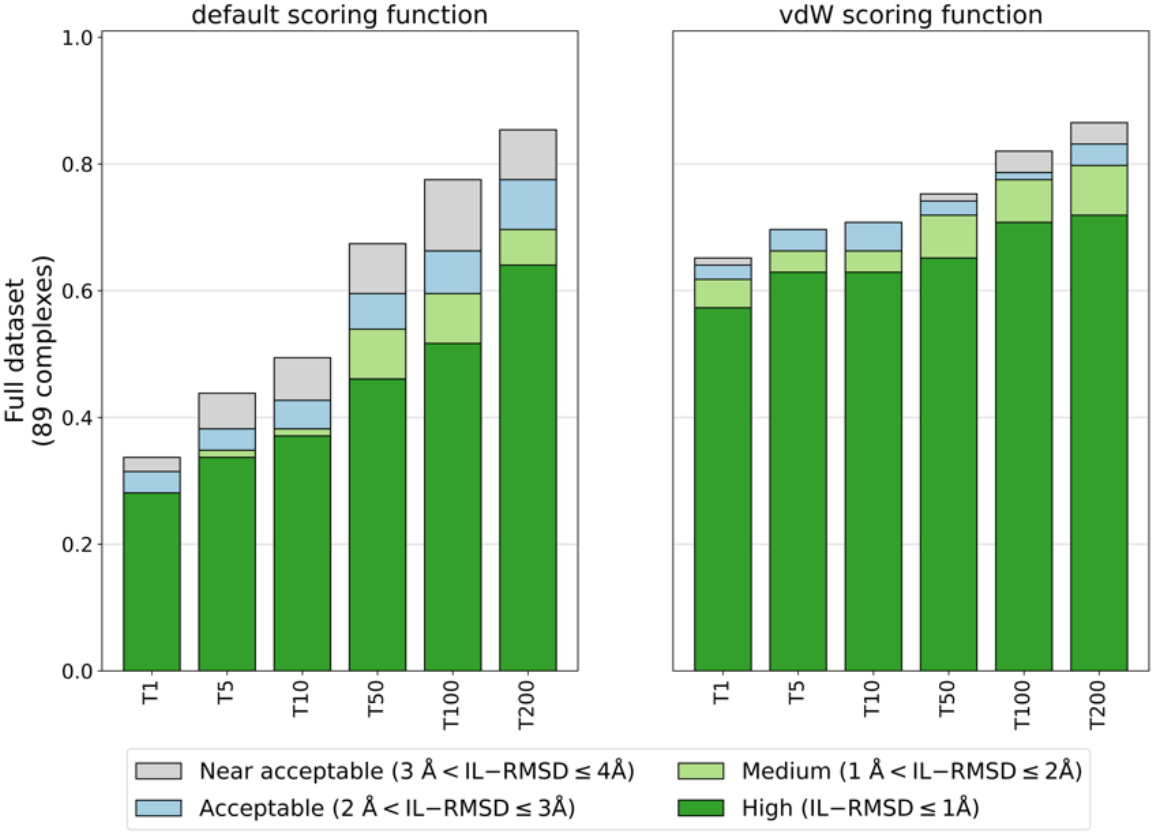
Comparison of bound docking success rates obtained with the *default* (w_vdW = 0.01) and *vdW* (w_vdW = 1.0) scoring functions (Eq. 1 and Eq. 2) as a function of the number of top-ranked models (T=1,5,10,50,100,200) selected, using true interface residues of both protein and glycan to define the ambiguous interactions restraints (**ti-aa** AIRs).

The *vdW* scoring function with increased van der Waals energy weight (w_vdW = 1.0) performs much better than the *default* one (w_vdW = 0.01) with remarkably higher success rates. This highlights the importance of the van der Waals energy term in scoring protein-glycans models. For example, considering T1 and T5 high-quality models, the improvement is around 30%. Even when considering the top200 models (the default number of models passed to the flexible refinement stage in a standard HADDOCK2.X workflow), a slight improvement is observed in both number and quality of acceptable of better models. Results are consistent if we consider the cluster-based success rate (see Methods), with 66% of top ranked clusters falling into the acceptable category in the *vdW* settings, compared to 33% of the *default* scenario. Glycans, despite all their polar groups, have quite hydrophobic properties, especially in their ring structure. A similar behaviour was observed in previous work when docking cyclic peptides^33^ and small ligands^27^. Based on these results, all subsequent docking calculations were performed with the *vdW* scoring function.

### Impact of glycan structural features and AIRs definition on the bound docking performance

The dependency of the structural features of the glycans on the SR was then assessed using the true-interface AIR restraints (**ti-aa**) and *vdW* scoring function scenario. SR were calculated grouping the complexes based on the size of the glycans (glycans composed by three or less units, labelled with **S**, or more than three, with **L**) and connectivity (linear, **L**, or branched, **B**). The SR, shown in the first column of <u>Supplementary Figure S2</u>, indicate that HADDOCK3 performs better for long linear (**LL**) and long branched (**LB**) glycans. The bound SR for T1 high quality models is 45%, 17%, 74%, and 73% for **SL, SB, LL**, and **LB** glycans, respectively (50%, 33%, 79%, and 82%, respectively, for T1 acceptable or better-quality models). When considering a larger number of models, the SRs become rather similar for the various types of glycans (70%-80% for T50 and 80-90% for T200), except for the short branched (**SB)** ones (∼ 70%). The lower performance on the short branched glycans, and, to a less extent on the **SL** ones, could be due to the fact that smaller ligands could be accommodated into the protein binding site with a greater variability of positions and orientations while the range of possible docking orientations is more restricted for larger ligands.

The impact of defining the glycan as passive (**tip-ap**) was then investigated. A slight decrease in the SR can be observed (second column of <u>Supplementary Figure S2</u>), with respect to the **ti-aa** scenario, for **LB** glycans and, to a less extent, for **SB** ones. This can be explained by noting that most of the linear long (**LL**) glycans (27/34) have all their residues involved in the binding to the protein, whereas the same is true for only half (5/11) of the long branched (**LB**) ones. Branched glycans are thus more difficult to model when no information about their interface is available. Overall, HADDOCK3’ performance on protein-glycan complexes is not much affected by defining the glycan as passive (**tip-ap** AIRs). As experimental interface information for the glycans might not always be available in a realistic scenario, the remaining of the manuscript will only discuss docking calculations with **tip-ap** AIRs. Note that, for example, Nuclear Magnetic Resource (NMR) can provide specific information about which groups of a glycan are involved in the binding, as demonstrated for the modelling of the complex between sialic acid and the N-terminal domain of the SARS-Cov2 spike protein using NMR saturation transfer experiments^34^.

### Unbound docking

In a realistic scenario, the bound conformations of the docking partners are unknown and only unbound structures or models will be available. As such, conformational rearrangements might be required during binding. We therefore assessed HADDOCK3’ performance on the *unbound* dataset consisting of 55 complexes. The starting point for the docking was the unbound form of the protein taken from the PDB and models of the glycans obtained with the GLYCAM-Web webserver^22,23^ (see Methods section *Dataset preparation*). When dealing with unbound structures, the flexible refinement stage of the workflow might allow some conformational changes to occur. As only a fraction (typically around 20%) of the models is subjected to flexible refinement, ranking and selection of models after the rigid-body docking stage becomes crucial. We tested two scenarios: 1) the “classical” scenario in which the top 200 ranked models are passed to the flexible refinement stage; 2) a cluster-based selection, made possible by the modular and flexible structure of HADDOCK3. In the second scenario, the rigid-body models are first clustered based on their RMSD similarity; then, the top 5 models of all the 50 clusters (150 when the ensemble is used, see below) are selected. In this way, models that might not rank high enough to be selected in an individual model ranking might still get selected and subjected to the flexible refinement, thus increasing the diversity of refined models (potentially at the cost of decreasing the overall number of acceptable or better models in the final set of refined models, which is not an issue provided the scoring function can identify the near native models). Since we are using an agglomerative hierarchical clustering method allowing to define the desired number of clusters, clusters with less than 5 models might be obtained and for this reason a different number of models might be subjected to the flexible refinement for each complex. The overall docking performance and the performance by glycan size and branching are shown in Figure 3.

**Figure 3.**
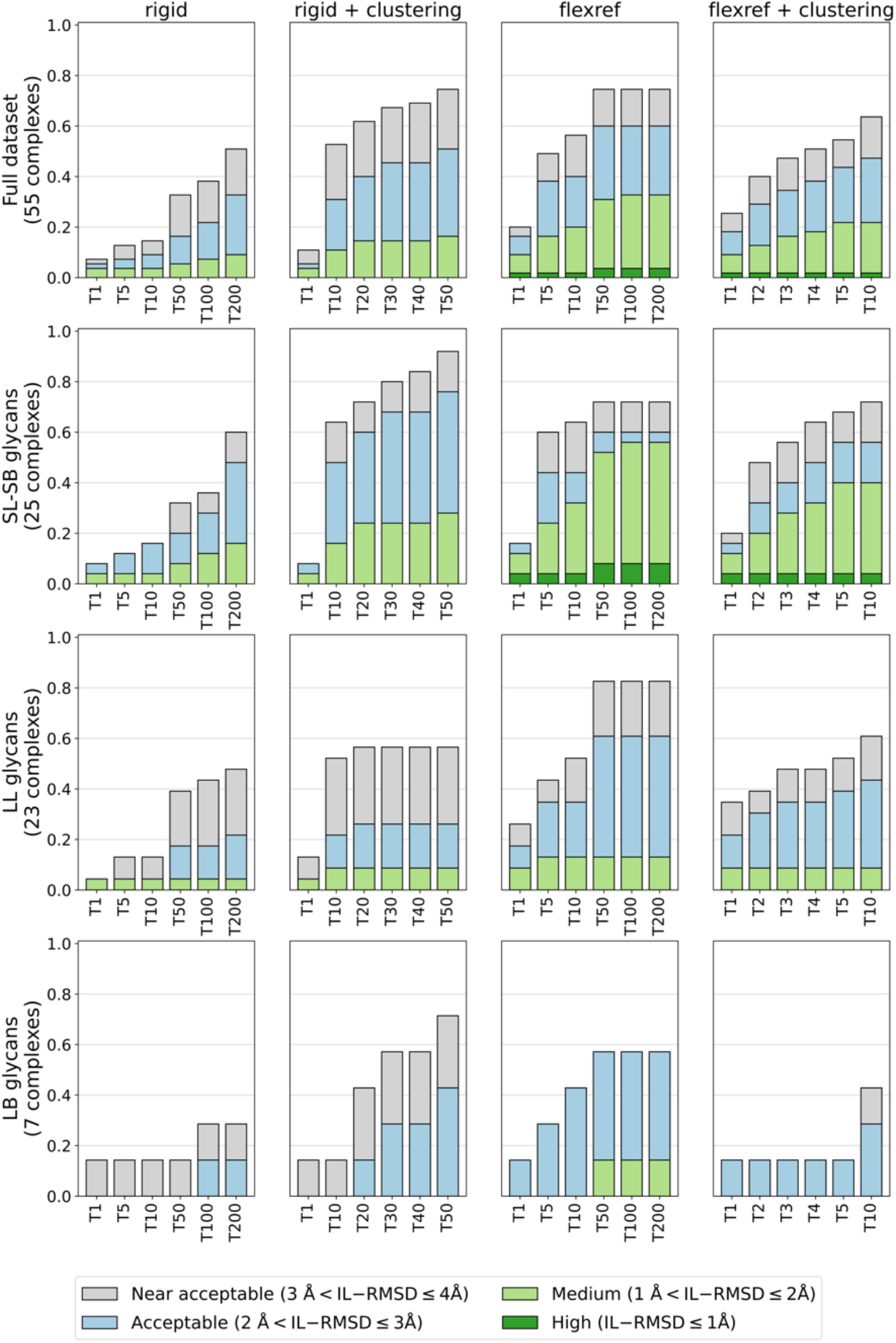
HADDOCK3 performance on the *unbound dataset*, using *vdW* scoring function and **tip-ap** AIRs (true interface on the protein defined as active and the glycan residues a passive). The success rates (SR) (Y axis), defined as the percentage of complexes for which an acceptable, medium or high quality models are generated, are calculated on the top 1 (T1) to top 200 (T200) ranked rigid body models (column “rigid”), T1 to T50 rigid body clusters, considering the top 5 models of each cluster (column “rigid + clustering”), the T1 to T200 ranked refined models (column “flexref”) and the T1 to T10 refined clusters, considering the top 4 models of each cluster (column “flexref + clustering”). SR are shown separately for the whole dataset (first row), and for the three categories of complexes grouped by glycans size and connectivity: **SL-SB** (second row), **LL** (third row), and **LB** (fourth row).

The comparison of the rigid body SR obtained on the single models (column “rigid” in Figure 3) with the SR obtained on the clustered models (column “rigid + clustering” in Figure 3), reveals that, overall, the selection of clustered models is beneficial to the docking success rate. This is particularly helpful for the long branched (**LB**) glycans (fourth row of Figure 3) with the clustering of the rigid body models allowing to retrieve more than 40% acceptable-quality models (70% near-acceptable-quality) compared to 15% acceptable-quality models (30% near-acceptable-quality) for the single model selection. Overall, for all types of glycans, selecting the rigid body models after clustering is the best way to choose the structures for the refinement stage.

The introduction of flexibility at the interface region strongly improves the quality of the models. Comparing the T200 refined models (column “flexref” in Figure 3) with the selected T50 rigid body clusters (column “rigid + clustering”), for short (**SL-SB**) glycans, almost 10% high-quality models are obtained, while there were none before, and almost 60% of the models falls within the medium quality cut-off, to be compared with around 30% at the rigid body stage. For the long linear (**LL**) glycans, the acceptable-quality success rate increases from around 25% (rigid body stage) to 61%. The improvement is also substantial for the **LB** glycans, for which the flexible refinement allows to obtain medium-quality models, which were not present at the rigid body stage, and an increase in acceptable-quality SR from 43% to 57%.

Finally, clustering of the refined models (fourth column in Figure 3) slightly improves the success rate compared to T10 single refined models, with around 50% of the glycans having an acceptable or better model in the top 10 clusters.

To demonstrate how flexible refinement affects both glycans’ conformations and models’ ranking, the best scoring refined models of three representative complexes are shown in Figure 4, superimposed onto their corresponding rigid body models and reference structures. These are representative of the **SL-SB, LL**, and **LB** groups. The refinement stage results in both better ranking and quality (lower IL-RMSD values) of the models. Overall, longer glycans are more challenging to refine than short ones. A high-quality model is produced for the 1C1L complex (**SL**, IL-RMSD = 0.52 Å after refinement), an acceptable-quality model for 5VX5 (**LL**, IL-RMSD = 2.22 Å), while for 1OH4 (**LB**, IL-RMSD = 4.42 Å), the quality is still not acceptable although it improved after the refinement stage.

**Figure 4.**
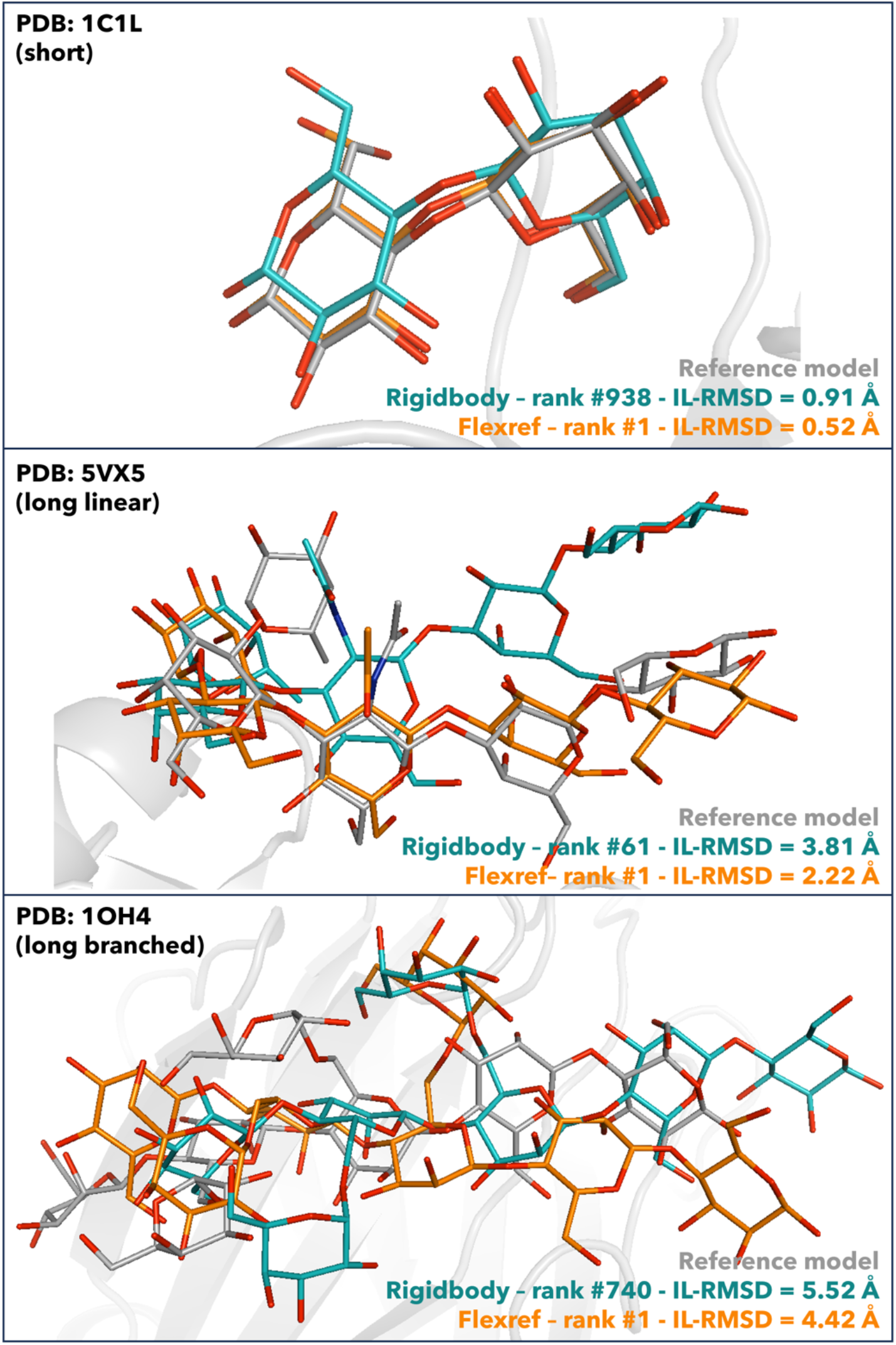
Superimposition of the best scoring flexible refinement models (orange) and the rigid body models (teal) to the reference structures (grey) for the complexes 1OH4 (**LB**), 5VX5 (**LL**), and 1C1L (**SL**) and the unbound docking scenario carried out with *vdW* scoring potential and **tip-ap** AIRs. Oxygen atoms of the glycans are shown in red in all the structures, nitrogens in blue, hydrogens not shown. Ranking and IL-RMSD values with respect to the reference structures for the flexref and rigidbody models are shown too.

We assess the overall impact of the refinement by measuring how much closer to the target structure is the best refined model with respect to the best model prior to the refinement stage: this analysis shows an average improvement of 0.43 Å on our dataset with a maximum observed improvement of 3.73 Å. Of course, not all models improve after refinement, but those that do improve also tend to be ranked higher by the HADDOCK scoring function, as shown in Fig.3. Proteins containing long linear glycans show the most substantial improvement (1.0 Å on average, with a maximum of 3.02 Å). Short (SL-SB) and long branched (LB) glycans show a negligible improvement (0.03 and 0.02 Å on average, respectively), although this is dependent on the specific glycan. For example, considering SL-SB glycans, we observe a maximum improvement of 3.73 Å (PDB 3AOF), while for some other models the flexible refinement worsens their quality. For LB glycans a maximum improvement of 2.13 Å is obtained (3AP9). Overall, the flexible refinement impacts most LL glycans.

We investigated whether increasing the length of the flexible refinement protocol would further improve the glycan conformations. This did not improve the success rates significantly nor in a uniform way, as it seemed to depend on the number of models passed to the refinement and the group of glycans considered (data not shown). It further comes at increased computational costs. As this behaviour could not be rationalized in a simple way, this approach was discarded.

### Can an ensemble of pre-sampled glycans conformations improve the docking performance?

While the performance in the *unbound docking* scenario is already quite high, unsurprisingly it does not reach the *bound docking* performance. One limiting factor here could be the unbound glycans conformations used for docking. Comparison of the conformations generated by the GLYCAM-Web webserver with respect to the bound ones reveals a strong dependency of the RMSD to the bound form on both the glycan’s size and connectivity with mean RMSDs increasing from 0.89Å for short linear glycans to 1.74 Å for long branched ones (see <u>Supplementary Figure S3</u>).

As HADDOCK can take an ensemble of conformations as starting point for the docking, we investigated if sampling conformations prior to docking could improve the overall docking performance. We used for that the water refinement module of HADDOCK varying the number of models generated and the length of the molecular dynamics sampling. Six different protocols were investigated (<u>Table S4</u>). The RMSD distributions from the bound glycan conformation of the various protocols are shown as box plots in <u>Supplementary Figure S4</u> and compared to the distribution of the original GLYCAM server conformations. From this analysis, the protocol that generates conformations closest to the bound form consists of sampling 400 models, with 16 times longer refinement protocol (sf400-x16) (which still remains a very short refinement protocol). With this sampling scenario, the average RMSD to the bound conformation decreases from 0.93 Å to 0.54 Å for **SL-SB** glycans, from 1.65 Å to 1.26 Å for **LL** glycans, and from 1.70 Å to 1.25 Å for **LB** glycans.

For docking, we want to limit the number of models in an ensemble to allow for sufficient sampling of each starting model combinations without having to increase the sampling too much. To this end we clustered the sampled glycan conformations using the `rmsdmatrix` and `clustrmsd` modules in HADDOCK3, requesting either 10 or 20 clusters from the ensemble of conformations. RMSDs from the bound conformation were calculated for the centres of each cluster with the `rmsdmatrix` module. Requesting 20 clusters results in the best sampling of the overall glycan RMSD distribution of the ensemble of 400 models, retaining low RMSD conformations. The cluster centres however rarely correspond to the best RMSD sampled, which results in some rather limited loss in RMSD to the bound form (see <u>Supplementary Figure S5</u>). Example of such conformational ensemble for the same glycans reported in Figure 4 are shown in <u>Supplementary Figure S6</u>.

The centres of those 20 clusters were provided as an ensemble for unbound docking, following the same rigid-body, cluster-based workflow described above. Only limited improvements are observed in single structure ranking performance after flexible refinement compared to that of the single unbound conformer protocol (see <u>Supplementary Figure S7</u>). For example, a higher number of medium-quality models for long linear glycans are obtained, together with more high-quality models for SL-SB glycans. No improvement is observed for long branched glycans. This can be attributed to the rather limited conformational sampling during the still short water refinement protocol (see <u>Supplementary Figure S6</u>). Better sampling strategies, possibly based on more extensive MD simulations will be required. Another issue is related to the selection of the relevant representative conformations, while limiting their number for docking purposes.

## Conclusions

In this study, we have assessed HADDOCK’s ability in modelling the 3D structures of protein-glycan complexes. First, the baseline performance was evaluated on the rather simple, unrealistic scenario starting from the partners in their *bound* conformations and giving full information about the interface (**ti-aa** AIRs). This allowed us to improve the rigid-body scoring function for protein-glycan complexes by increasing the weight of the intermolecular van der Waals energy term to 1.0, as done for the docking of small molecules to proteins^27^. An analysis of the *bound docking* performance per types of the glycans revealed that longer glycans (in their bound conformation) are easier to model. This is probably a consequence of the lower number of possible orientations that longer glycans can assume when binding the protein. In the more realistic *unbound docking* scenario, in which the glycans conformations were modelled with the GLYCAM-Web webserver and free form structures of the proteins are used, conformational changes are required to generate native-like poses. In this case, longer glycans are more difficult to model with only (near-) acceptable predictions generated in most cases.

Making use of the new modular version of HADDOCK3, we introduced a protocol in which rigid-body models are clustered prior to refinement, with representatives of all clusters being passed to the flexible refinement stage of HADDOCK. This strategy enables HADDOCK to retain 51% acceptable or better models after rigid-body, compared to 33% using the standard single model selection. After flexible refinement, the overall performance reaches an overall success rate of 16% (resp. 18%) and 38% (resp. 44%) for T1 and T5 single structure models (resp. clusters).

Single structure or cluster-based selection of models at the end of the workflow show almost similar performance. This is in line we what was also observed in small molecule docking with HADDOCK^27,35^.

While the flexible refinement does improve the quality of the models, it is not sufficient in cases where large conformational changes are required. We therefore investigated if a limited sampling of glycan conformations prior to docking using the water refinement module of HADDOCK could generate conformations closer to the bound form. While we do observe slight improvements, their impact on the docking performance in an ensemble docking scenario is limited, thus highlighting the need for more extensive conformational sampling strategies, together with methods to identify the most relevant conformations. For example, the use of a database of glycans conformations such as GlycoShape^36^, could help in improving HADDOCK’s performance in modelling protein-glycan complexes.

## Supporting Information

▪ *Supplementary_Table_S1*.*xlsx* includes all the information about the dataset used in the study
▪ *Supplementary_Information*.*pdf* includes all supplementary texts, figures and tables.

## AUTHOR INFORMATION

### Author Contributions

The manuscript was written through contributions of all authors. All authors have given approval to the final version of the manuscript.

## ABBREVIATIONS

HADDOCK: High Ambiguity Driven DOCKing
IL-RMSD: Interface-Ligand Root Mean Squared Deviation
SL: short-linear
SB: short-branched
LL: long-linear
LB: long-branched
AIRs: Ambiguous Interaction Restraints
PDB: Protein Data Bank.

