## Supplementary Information for "Modelling Protein-Glycan Interactions with HADDOCK"

**Table of Contents**


- Text S1. Details on the preparation of proteins and glycans structures……..……….…p. S2
- Text S2. Details on glycans conformational sampling……………………..…………...p. S2
- Table S2. Modules and parameters used for bound docking……………………..…… p. S3
- Table S3. Modules and parameters used for unbound docking………………..………p. S4
- Table S4. Glycans conformational sampling scenarios…………………..……………p. S5
- Figure S1. Example of HADDOCK models satisfying the quality thresholds……......p. S6
- Figure S2. HADDOCK3 performance on the bound dataset………………………… p. S7
- Figure S3. Glycans RMSD to their bound conformations…………………………… p. S8
- Figure S4. Impact of mdref on glycans conformations……………………………… p. S9
- Figure S5. Impact of the clustering on glycans lowest RMSD……………………… p. S10
- Figure S6. Examples of glycans ensembles of conformations……………………… p. S11
- Figure S7. HADDOCK3 performance with the ensembles of glycans………………p. S12
- Supplementary References…………………………...…………………….…….. p. S13

**Text S1: Details of the preparation of proteins and glycans structures**

Glycans unbound conformations were generated with the GLYCAM-Web webserver[1,2] with the GLYCAM nomenclature for the carbohydrates residues modified to match the naming recognized by HADDOCK. Prior to docking calculations, all structures were pre-processed with a combination of pdb-tools[3], in-house scripts and manual curation. Heteroatoms such as water molecules, cofactors and ions were removed when not part of the protein – glycan interface. The residues were renumbered to start from 1, with `pdb_reres`, and the chains ID were modified using `pdb_chain` to chain A and chain B for the receptor and the ligand, respectively. In some cases where the receptor consists of more than one chain, these were merged into a single chain, and the residue numbering was shifted if needed to avoid overlap in numbering. Alternative occupancies were removed by selecting the conformation with the highest occupancy value.

**Text S2: Details of the glycans conformational sampling protocol**

Glycans conformations generated from the GLYCAM-Web webserver were used as starting points for conformational sampling. For 11 out of 55 glycans two conformations were generated by the webserver (four for 1OH4); for those cases, all conformations were used. Conformational sampling and analysis were performed with HADDOCK3 using the following modules: `topoaa, mdref, rmsdmatrix, clustrmsd`. After the creation of the topology (module `topoaa`), conformational sampling in water was performed with the water refinement module (`mdref`), defining the glycans as fully flexible (parameters: `nfle1 = 1; fle_sta_1_1 = 1; fle_end_1_1 = 7`). At this stage, different scenarios were tested in terms of number of steps and number of models, as specified in Table S4. Three scenarios were run on 100 models with increasing number of steps / simulation time (sf100-x1, sf100-x8, sf100-x16). Then, the number of models was increased to 400 while the simulation time was the same of two of the scenarios previously listed (sf400-x1 and sf400-x16). The overall time of the simulations ranges from 185000 (sf100-x1) to 11240000 steps (sf400-x16).
For assessing the conformational variability over the sampled trajectory, all-atom RMSD were calculated for all the generated conformations, with respect to the bound conformations, with the HADDOCK3 `rmsdmatrix` module. RMSD distributions of the conformations obtained with the sampling procedures were plotted together with the RMSD calculated for the webserver-generated conformations with respect to the bound conformations.

The RMSD matrix between all the conformations generated was calculated with the `rmsdmatrix` module, by specifying, through the parameter '`resdic_`', the residues to be considered for the alignment and the RMSD calculation. The `clustrmsd` module was then exploited for clustering the conformations, with the following parameters: `criterion = maxclust, linkage = average, n_clusters = 10 (or 20)`. The '`maxclust`' criterion clusters the structure in such a way to give a fixed number of clusters, defined by the parameter `n_clusters`. The linkage governs the way clusters are merged in the creation of the dendrogram, i.e., it defines the method for calculating the distance between the newly formed cluster and each object which does not belong to a cluster yet. RMSD with respect to the bound conformations were calculated for the clusters centers, i.e. the points having the lower distance to all the other points in the cluster. RMSD of to the cluster centers were plotted together with the overall sampling distribution to assess whether the clustering can capture the models that are closer to the glycan experimental structure. The centers of the clusters were then used as an ensemble for further docking calculations.

**Table S2**. Modules and parameters used for bound docking.

| **Protocol bound dataset** | | |
|---|---|---|
| **Stage** | **Module** | **Parameters** |
| 1 | `topoaa` | |
| 2 | `rigidbody` | `sampling = 1000`<br>`w_vdw= 0.01 (default), 1.0 (vdW)`<br>`ambig_fname = /path/to/tbl/file` |
| 3 | `caprieval` | `reference_fname = /path/to/reference/pdb` |
| 4 | `rmsdmatrix` | `resdic_A = [interface residues of the protein]`<br>`resdic_B = [interface (`**`ti-aa`**`) or all (`**`tip-ap`**`) residues of the glycan]` |
| 5 | `clustrmsd` | `criterion = distance`<br>`linkage = average`<br>`min_population = 4`<br>`clust_cutoff = 2.5 Å` |
| 6 | `caprieval` | `reference_fname = /path/to/reference/pdb` |

**Table S3.** Modules and parameters used for *unbound docking*.

| **Protocol unbound dataset** | | |
|---|---|---|
| **Stage** | **Module** | **Parameters** |
| 1 | topoaa | |
| 2 | rigidbody | sampling = 1000, 4000 (ensemble)<br>w_vdw= 1.0 (vdW)<br>ambig_fname = /path/to/tbl/file |
| 3 | caprieval | reference_fname = /path/to/reference/pdb |
| 4 | rmsdmatrix | resdic_A = [interface residues of the protein]<br>resdic_B = [all residues of the glycan] |
| 5 | clustrmsd | criterion = maxclust<br>n_clusters = 50, 150 (ensemble) |
| 6 | seletopclusts | top_models = 5 |
| 7 | caprieval | reference_fname = /path/to/reference/pdb |
| 8 | flexref | tolerance = 5<br>nemsteps = 200<br>mdsteps_rigid = 500<br>mdsteps_cool1 = 500<br>mdsteps_cool2 = 1000<br>mdsteps_cool3 = 1000<br>ambig_fname = /path/to/tbl/file |
| 9 | caprieval | reference_fname = /path/to/reference/pdb |
| 10 | rmsdmatrix | resdic_A = [interface residues of the protein]<br>resdic_B = [all residues of the glycan] |
| 11 | clustrmsd | criterion = distance<br>linkage = average<br>min_population = 4<br>clust_cutoff = 2.5 Å |
| 12 | caprieval | reference_fname = /path/to/reference/pdb |

**Table S4.** Glycans conformational sampling scenarios.

| scenario_name | Sampling_factor | waterheatsteps | watersteps | watercoolsteps | Total number of steps |
|---|---|---|---|---|---|
| Default mdref values | 1 | 100 | 1250 | 500 | |
| sf100-x1 | 100 | 100 | 1250 | 500 | 185000 |
| sf100-x8 | 100 | 100 | 10000 | 4000 | 1410000 |
| sf100-x16 | 100 | 100 | 20000 | 8000 | 2810000 |
| sf400-x1 | 400 | 100 | 1250 | 500 | 740000 |
| sf400-x16 | 400 | 100 | 20000 | 8000 | 11240000 |

Changes with respect to the scenario sf100-x1 are highlighted in blue. Scenario sf100-x1 follows default parameter setting except for the sampling factor (i.e. number of models generated). The last column reports the total number of steps defined as: sampling_factor * (waterheatsteps + watersteps + watercoolsteps).

**Figure S1**. Example of HADDOCK models for three different PDB files satisfying the four different quality thresholds highlighted in the main text, namely high, medium, acceptable and near acceptable. Short glycans are clearly more sensitive to small deviations, thus making the near acceptable (IL-RMSD between 3.0 Å and 4.0 Å) threshold too lenient. For long linear glycans (2ZEX in the figure), near acceptable models can still be useful for downstream analysis.

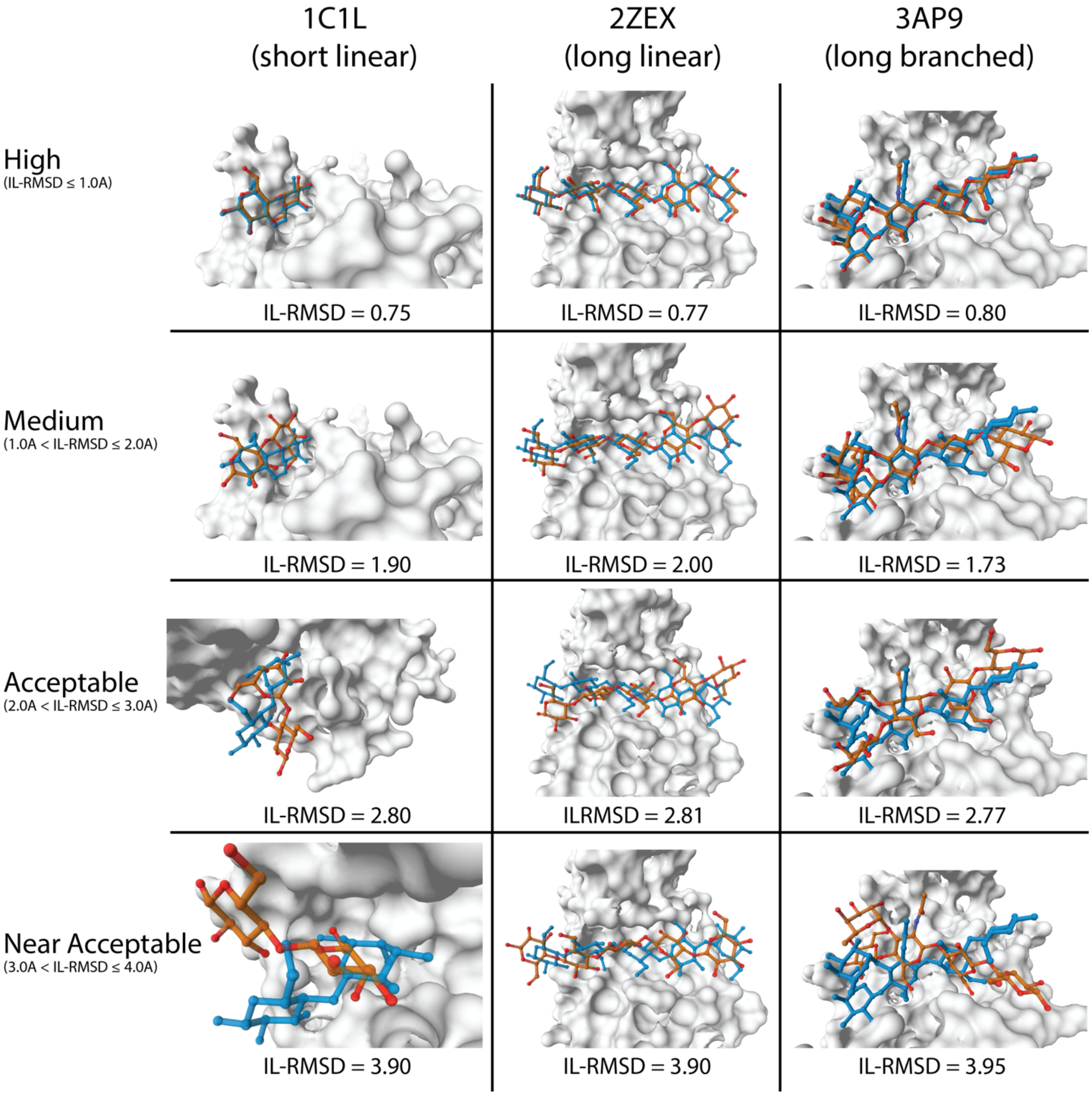

**Figure S2**. HADDOCK3 performance on the *bound dataset*, as a function of glycans size (**S**: glycans composed by three or less monosaccharide units, or **L** by more than three) and connectivity (**L**: linear, or **B**: branched). The left column shows the performance using both the protein and glycan interface residues as active (**ti-aa**) and the right column using the protein interface residues as active and the entire glycan as passive (**tip-ap**). Success rates are calculated for the top (T) 1, 5, 10, 50, 100, and 200 models.

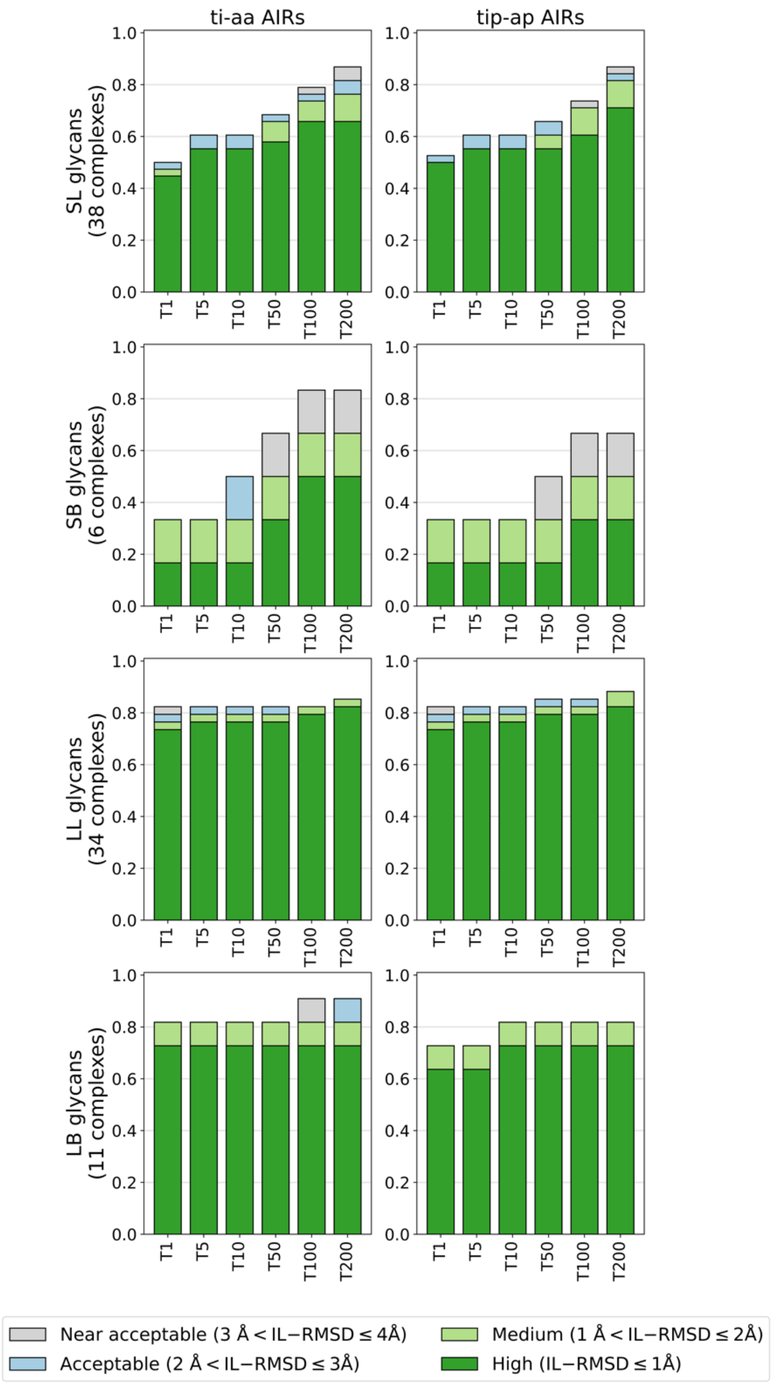

**Figure S3.** Violin plot showing the distribution of glycan RMSD between the conformations generated with the GLYCAM server and the corresponding bound structure for the three categories of glycans **SL-SB**, **LL**, and **LB**. Mean, maximum and minimum values are indicated in the plot.

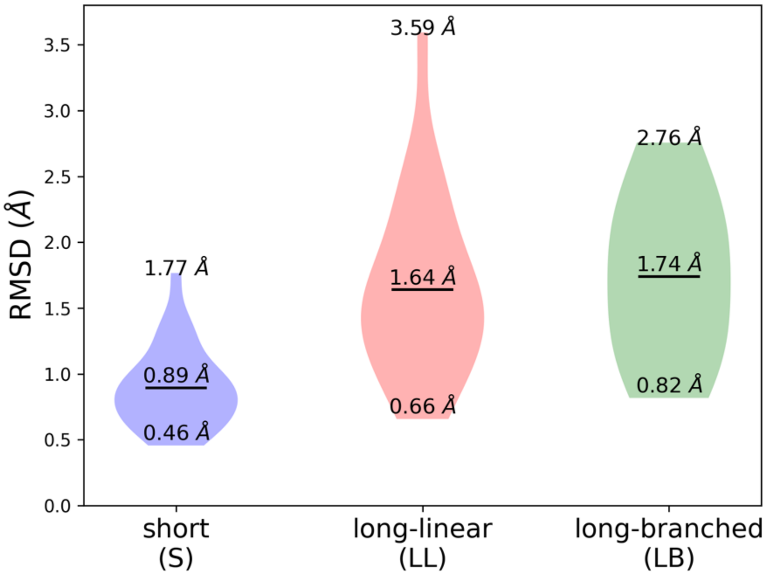

**Figure S4.** Boxplots of the reference glycan RMSD distribution of the GLYCAM server-generated conformations (left) and the lowest RMSD to the bound form obtained with the 5 sampling scenarios. The comparison is shown for the three groups of glycans: **SL-SB** (top), **LL** (middle), and **LB** (bottom).

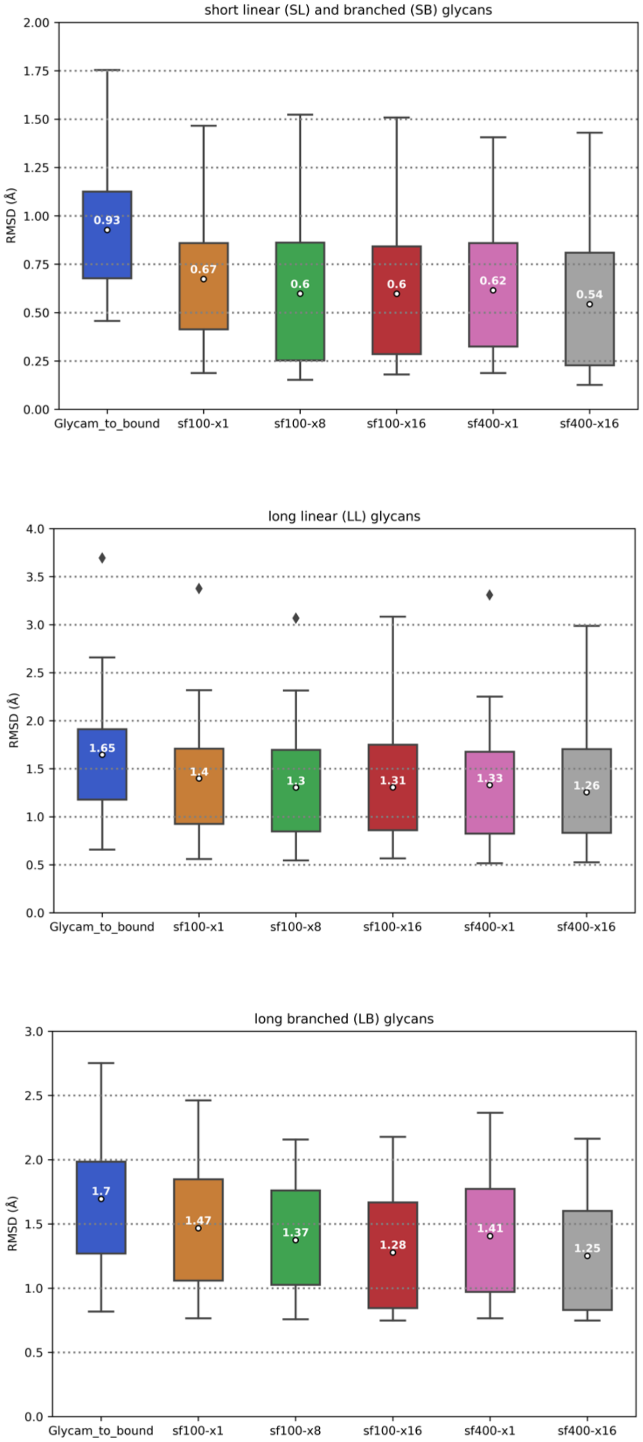

**Figure S5.** Comparison between lowest RMSDs (glycans sampled conformations with respect to the bound forms) after clustering vs lowest RMSDs from the overall sampling. The comparison is shown for the three groups of glycans: short (**SL-SB**, top left), long linear (**LL**, top right), and long branched (**LB**, bottom).

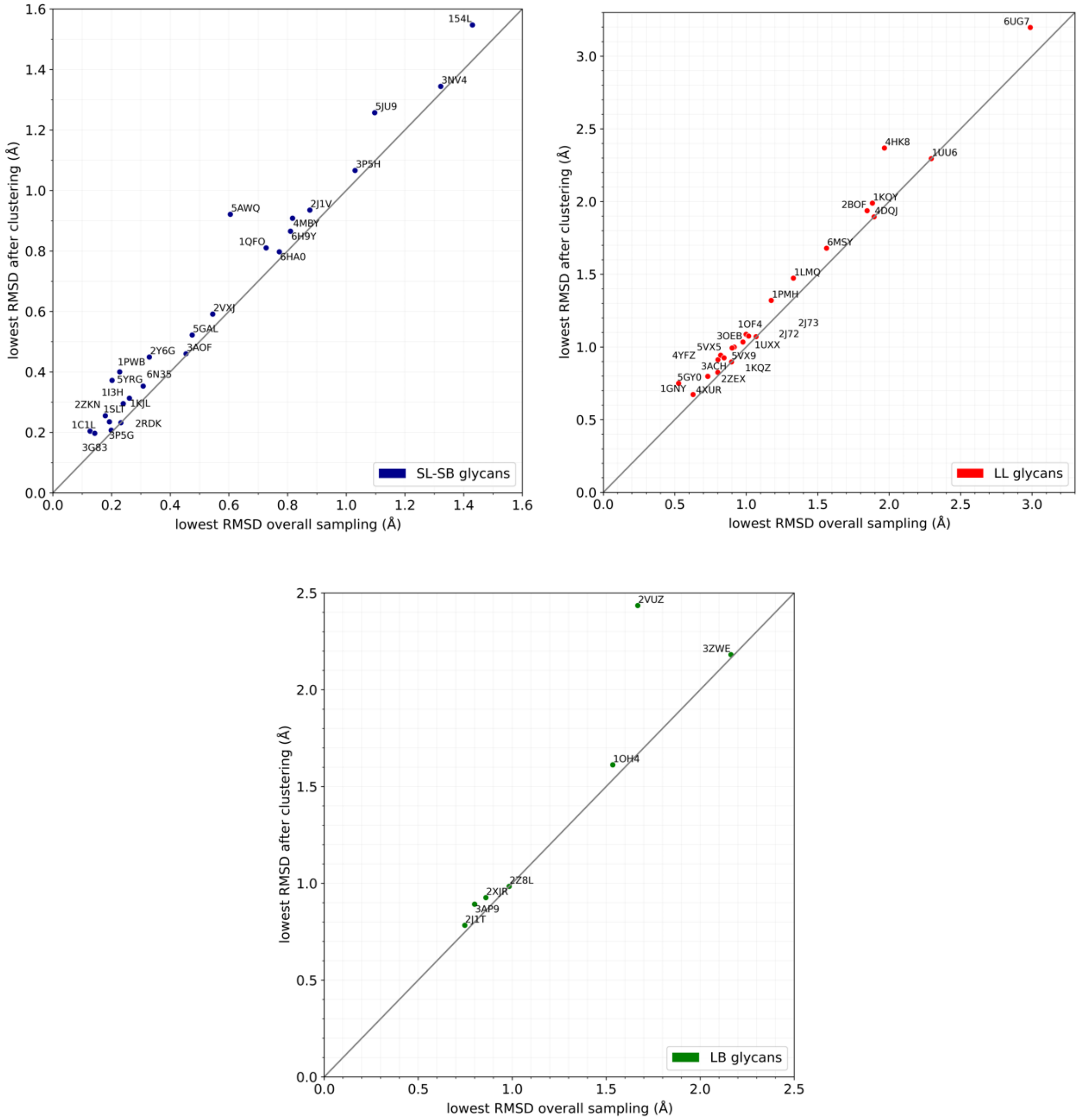

**Figure S6.** Superimposition of the centers of the 20 clusters (carbon atoms in light blue) obtained from the sf400-x16 sampling scenario and of the unbound conformation generated by GLYCAM-Web webserver (carbon atoms in yellow) to the bound conformations (carbon atoms in black) for the for the complexes 1OH4 (**LB**), 5VX5 (**LL**), and 1C1L (**SL**). Oxygen atoms are shown in red in all the structures, nitrogens in blue, hydrogens not shown.

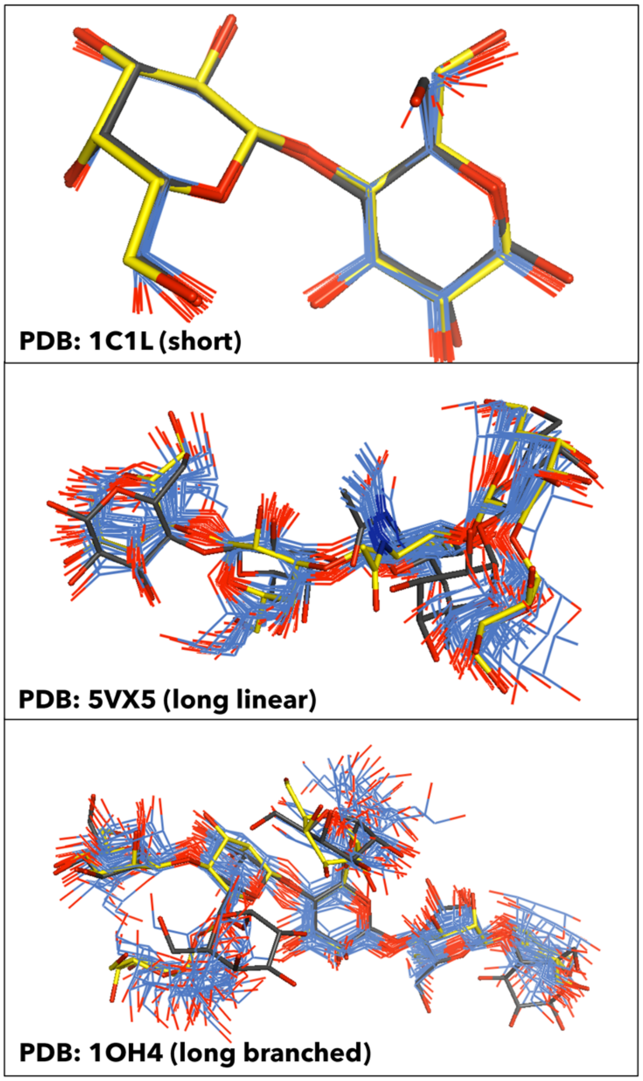

**Figure S7.** HADDOCK3 performance on the *unbound dataset*, using *vdW* scoring function and **tip-ap** AIRs. The success rates (SR), calculated on the top (T) 1, 5, 10, 50, 100, and 200 refined models (flexref stage), are compared between single conformations runs (left column) and ensemble runs (right column). SR are calculated for the entire dataset (first row), and separately for the three categories of complexes grouped by glycans size and connectivity: **SL-SB** (second row), **LL** (third row), and **LB** (fourth row).

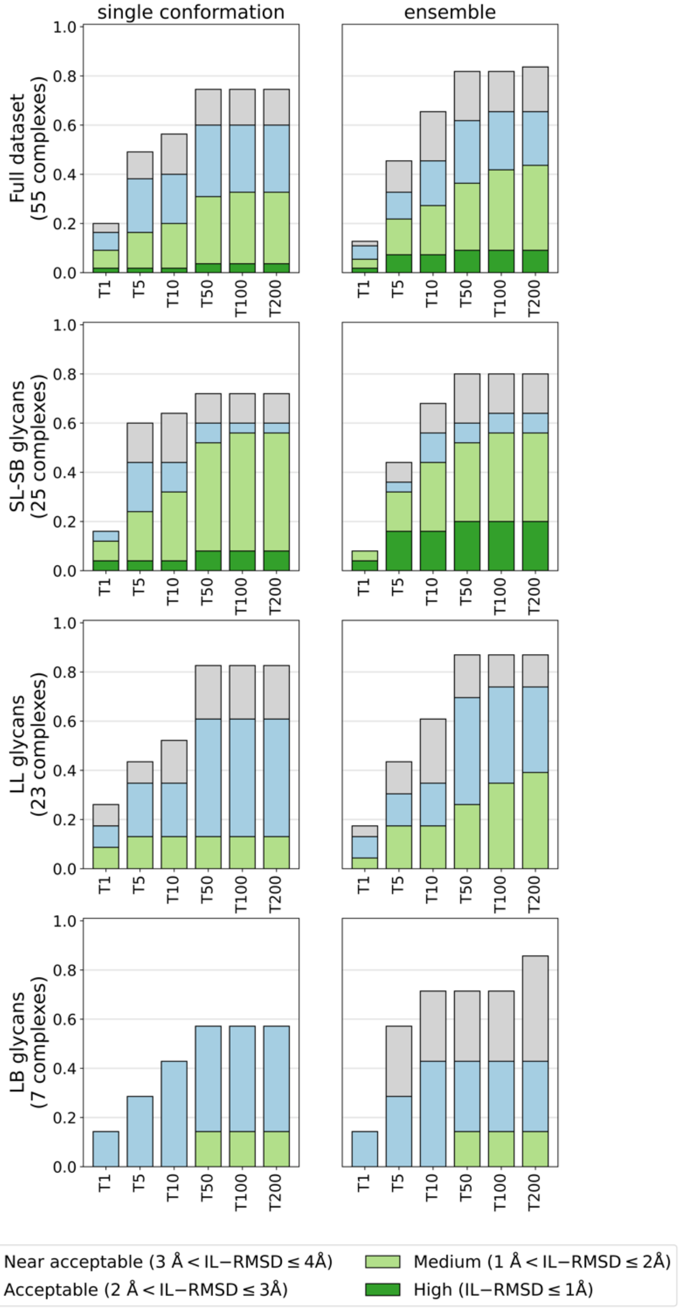
